# Females with Obesity Exhibit Greater Influenza Vaccine-induced Immunity and Protection than Males in a Mouse Model

**DOI:** 10.1101/2025.11.18.689040

**Authors:** Brian Wolfe, Saurav Pantha, Saranya Vijayakumar, Shristy Budha Magar, Tawfik Aboellail, Santosh Dhakal

**Author notes:** Correspondence: Dr. Santosh Dhakal, Department of Diagnostic Medicine/Pathobiology (DMP) Kansas State University (KSU), Manhattan, KS, 66506. The authors contributed equally to this project (Brian Wolfe, Saurav Pantha).

## Abstract

**Introduction:** Obesity is increasing globally, and it negatively impacts influenza vaccine efficacy. Although sex differences in influenza vaccine responses are studied in non-obese hosts, studies investigating sex differences in influenza vaccine-induced immunity and protection during obesity are limited.

**Materials and methods:** Using the C57BL/6J mouse model of high-fat diet (HFD)-induced obesity or low-fat diet controls, we investigated sex differences in influenza vaccine-induced immunity and protection during obesity. Male and female mice with or without obesity were vaccinated intramuscularly twice at a 3-week interval with an inactivated 2009 H1N1 influenza A virus (IAV) vaccine. At 35 days post-vaccination (dpv), antibody responses in plasma and B- and T-cell responses in spleen and bone marrow were quantified. At 42 dpv, mice were intranasally challenged with a drift variant of the H1N1 IAV, and disease severity was assessed by monitoring the change in body mass up to 21 days post-challenge (dpc). Subsets of mice were euthanized at 3 dpc to determine pulmonary virus replication (TCID_50_ assay), histopathology (H&E staining), and cytokine/chemokine responses (multiplex ELISA).

**Results:** Female mice, irrespective of diet and obesity status, developed higher antibody responses and were better protected compared to males. Vaccinated males with obesity mounted the poorest antibody responses, experienced a more severe disease, were unable to clear replicating virus from the lungs effectively, and demonstrated heightened pulmonary inflammation. Despite these differences, splenic B- and T-cell frequencies were comparable, suggesting the inefficiency of B cells to produce antibodies in males but not in females with obesity.

**Discussion:** Our findings suggest that sex differences is observed in influenza vaccine-induced immunity and protection during obesity, where males are more severely affected. These findings highlight the importance of considering biological sex and obesity status in influenza vaccine design and testing.

## Introduction

Obesity is an independent risk factor for influenza-associated disease severity and mortality (1, 2). Individuals with obesity shed the influenza virus for a longer duration than healthy-weight individuals, both when symptomatic and asymptomatic (3). They are also more likely to develop lower respiratory tract infections during influenza virus infection and require prolonged hospitalization (4). Therefore, vaccination is one of the major means of reducing the toll of influenza in the population with obesity.

Epidemiological studies show mixed results about the effects of obesity on influenza vaccine effectiveness. Some studies suggest similar vaccine effectiveness compared with healthy-weight populations (5, 6), whereas others demonstrate a more rapid decline in antibody titers (7), and a two-fold higher risk of influenza-like illnesses in the vaccinated adults with obesity compared to healthy weight adults (8). Differences in study population, season, vaccine strains, mismatches with circulating virus strains, study endpoints, and other confounding factors, such as comorbidities, may explain variations in epidemiological findings. Studies in the mouse model, which provides a more controlled environment, suggest that obesity is associated with a rapid decline in antibody responses and an inferior protective efficacy of influenza A virus (IAV) vaccine, which cannot be improved even after increasing the vaccine dose or following the addition of adjuvants (9, 10). These findings suggest that the correlates of protection against IAV infection may differ between individuals with or without obesity.

Human and mouse studies, predominantly carried out in non-obese hosts, have shown that biological sex (i.e., the differences between males and females based on sex chromosomes, gonadal tissues, and sex steroids) influences IAV pathogenesis and vaccine-induced immunity (11–13). In humans, following vaccination with seasonal or pandemic influenza vaccines, adult females of reproductive age develop a greater antibody response than males (14–17). A recent meta-analysis of phase 3 randomized trials (2010-2018) also showed higher immunogenicity and vaccine effectiveness in females than in males (18). Mouse model studies align with these observations: female mice produce around two times greater virus-specific antibodies, generate a greater number of germinal center (GC) B cells with superior somatic hypermutation (SHM) frequencies, and are better protected from IAV challenge than males (15, 17, 19). These differences are contributed to by sex hormones (e.g., testosterone and estradiol) and genetic factors such as higher expression of *Tlr7* gene in females (15, 17).

Despite these insights, studies investigating sex differences in influenza vaccine-induced immunity and protection in the context of obesity are limited. We hypothesized that sex differences in influenza vaccine responsiveness are maintained during obesity, with females exhibiting superior antibody responses and better protection. In this study, we used a high-fat diet-induced obesity mouse model to investigate how biological sex influences influenza vaccine-induced antibody responses and protection following immunization with an inactivated 2009 pandemic H1N1 IAV vaccine.

## Materials and methods

### Animals

C57BL/6J mice (4-5 weeks old; strain 000664) were purchased from the Jackson Laboratory (Bar Harbor, Maine, U.S.), and after a week of acclimatization, they were randomly assigned either to a low-fat diet (LFD;10%kCal, D12450J, Research Diets, New Brunswick, NJ, U.S.) or high-fat diet (HFD;60%kCal, D12492i, Research Diets, New Brunswick, NJ, U.S.) (20). After 12-weeks of diet treatment, the body mass of each mouse on HFD was compared with the average body mass of mice on LFD, and obesity was defined as a body mass of ≥20% greater than the average body mass of age- and sex-matched mice on LFD. Mice maintained on HFD for 12 weeks, but that did not reach the obesity threshold, were considered non-responders and excluded from the obesity study but analyzed separately as non-responder group. Animal procedures were carried out at Kansas State University (KSU) as per the Institutional Animal Care and Use Committee approval (protocol number 4855).

### Glucose tolerance test (GTT)

For GTT, after a 6-hour fast, mice received an intraperitoneal injection of 25% glucose solution at a dose of 2g/Kg body mass. Blood glucose levels were measured at 0, 30, and 120 minutes after glucose administration using the AlphaTrak3 blood glucose monitoring system (Zoetis, NJ, USA) by pricking the tip of the tail with a sterile lancet (20).

### Vaccination, challenge, and morbidity measurement

Mice were vaccinated intramuscularly with an inactivated H1N1 IAV vaccine (A/California/04/2009; 2009 pandemic strain; 20µg/40µL PBS) and boosted after 21 days (19, 21, 22). At 35 dpv, blood was collected to measure antibodies, and subsets of mice were euthanized to quantify B- and T-cell responses. At 42 dpv, mice were challenged intranasally under ketamine (80-100 mg/mL) and xylazine (5-10 mg/mL) anesthesia with 10^5^ tissue culture infectious dose 50% (TCID_50_) units of a drift variant of the 2009 pandemic H1N1 IAV strain (19, 21, 22). During the day of virus infection, when mice were anesthetized, their nose-to-anus length was also measured to calculate the body mass index (BMI; body mass/length²). Following virus challenge, body mass was recorded daily up to 21 days post-challenge (dpc) to monitor disease severity. Mice that lost more than 25% of the baseline body mass were humanely euthanized (22). Subsets of mice were euthanized at 3 dpc for the collection of lungs to compare virus titers (TCID_50_ assay), cytokine/chemokine responses, and histopathological changes. Mice immunized with PBS (vehicle only) but challenged with the same viral dose and euthanized at 3 dpc were used as controls.

### Antibody measurement by enzyme-linked immunosorbent assays (ELISAs)

In-house standardized ELISAs were used to measure IgG, IgG1, and IgG2c antibodies (19, 21, 22). Briefly, high-binding ELISA plates (Greiner Bio-One, NC) were coated with the whole virus protein of the 2009 H1N1 pandemic IAV (100ng/well) in sodium carbonate/bicarbonate buffer. A 10% skim milk solution (200µL/well) was used for blocking. Serially diluted heat-inactivated plasma samples (starting from a 1:250 dilution) were transferred in duplicates to the coated plates. Secondary antibodies used were IgG (#31430, Invitrogen, CA, USA), IgG1 (#PA1-74421, Invitrogen, CA, USA), and IgG2c (#56970, Cell Signaling Technologies, MA, USA), followed by the tetramethylbenzidine (TMB) substrate reagent (#555214, BD Biosciences, CA, USA) addition (50µL/well). To stop color development 1M Hydrochloric Acid (HCL, 50µL/well) was added. Absorbance was measured at 450nm on a spectrophotometer (Biotek, VT, USA).

### Microneutralization assay

Virus neutralization antibody (nAb) titers were measured using the Madin-Darby Canine Kidney (MDCK) cells infection method. Briefly, two-fold serially diluted plasma samples (beginning at 1:20 dilution) were incubated with 100 TCID_50_ of the 2009 pandemic H1N1 IAV for 1 hour at room temperature and transferred in duplicates (50µL/well) to the cell culture plates and incubated in 32^0^C in a 5% CO_2_ incubator. In 24 hours, plates were washed once with PBS and fresh medium was added. After an additional 6 days of incubation, cell plates were fixed using 4% formaldehyde solution and stained with naphthol blue black solution (19, 21, 22). Neutralizing titers were defined as the reciprocal of the highest plasma dilution that prevented virus-induced cytopathic effect.

### Virus titration in lung homogenates

Infectious virus titers in lung homogenates were quantified by tissue culture infectious dose 50% (TCID_50_) assay (19, 21, 22). Right lung lobes were homogenized in DMEM at a 1:4 weight/volume ratio, and clarified supernatants were diluted 10-fold (22). Dilutions were added in six replicate wells of 96-well plates containing confluent MDCK monolayers and incubated at 32°C in 5% CO_2_ incubator for 6 days. Following incubation, cells were fixed with 4% formaldehyde and stained with naphthol blue-black. Wells were scored for cytopathic effects, and virus titers were calculated using the Reed and Muench method as log_10_ TCID_50_/mL (19, 21, 22).

### Histopathology

Left lung lobes were inflated via trachea with zinc-buffered formalin (Z-Fix, Anatech, MI, USA) and immersed in fixative. Samples were processed by the Kansas Veterinary Diagnostic Laboratory, where lungs were paraffin-embedded, sectioned at 5 µm thickness, mounted on glass slides, and stained with hematoxylin and eosin (H&E). Lung sections were evaluated by a board-certified veterinary pathologist in a single-blinded manner. Pulmonary inflammation was scored on a semiquantitative 0-5 scale across the following parameters: pleural inflammation (max score 5); airway inflammation (main stem bronchus, primary bronchiole, and right bronchus, max total 15); parenchymal inflammation (max 5); and perivascular inflammation (arteirs, veins, and capillaries, max total 15). The maximum cumulative score was 40 (20).

### Cytokines/chemokines analysis

Cytokine and chemokine concentrations in lung homogenates were quantified using the ProcartaPlex mouse cytokine and chemokine panel 1, 26-plex assay (#EPX260-26088-901, Thermo Fisher Scientific, Waltham, MA, USA) as per the manufacturer’s instructions. Samples were run in singlets due to volume limitations, whereas standards were run in duplicates. Plates were read on a Luminex xMAP technology (Austin, TX, USA), concentrations (pg/mL) were interpolated from standard curves using the ProcartaPlex Analysis App (Thermo Fisher Scientific, Waltham, MA, USA) (22).

### Cell preparation for flow cytometry

Single-cell suspensions from spleens were prepared by pressing tissues through a 70μm cell strainer into ice-cold fluorescence-activated cell sorting (FACS) buffer (PBS supplemented with 25mM of HEPES and 1mM of EDTA) (21). Bone marrow cells were obtained by collecting the femur and tibia from the vaccinated side, cutting both ends of each bone, and flushing the marrow cavity with ice-cold FACS buffer (23). Red blood cells were lysed with ACK lysis buffer (#A1049210, Gibco, Waltham, MA, USA), and cells were washed, resuspended in FACS buffer, and passed through a 70 μm cell strainer before staining. Cell concentration and viability were determined by trypan blue exclusion method.

### Flow cytometry

For flow cytometry, 5 x 10^6^ splenocytes and 1 x 10^6^ bone marrow cells were used per animal. Two antibody panels were designed: panel 1 to identify germinal center (GC) B cells, plasmablasts, plasma cells, and memory B cells, and panel 2 to identify follicular T helper (Tfh) cells (19, 21). Cells were incubated with rat anti-mouse CD16/CD32 (#553142, clone: 2.4G2, BD Biosciences, CA, USA) to block non-specific binding of antibodies to Fc receptors and stained with fixable viability stain (FVS) 780 (#565388, BD Biosciences) to exclude dead cells. Antibodies used in panel 1 were: FITC rat anti-mouse CD4 (#561828, clone: GK1.5, BD Biosciences), PE-Cy7 rat anti-mouse CD45R/B220 (#561881, clone: RA3-6B2, BD Biosciences), BV421 rat anti-mouse CD38 (#562768, clone: 90/CD38, BD Biosciences), PE rat anti-mouse T- and B-cell activation antigen (GL7) (#561530, clone: GL7, BD Biosciences), APC rat anti-mouse CD138 (#561705, clone: 281-2, BD Biosciences), BV786 rat anti-mouse IgD (#563618, clone: 11-26c.2a, BD Biosciences), RB705 mouse anti-mouse IgM (#757936, clone: AF6-78, BD Biosciences), and PerCP-eFluor 710 anti-mouse IgD (#46599382, clone: 11-26c, eBioscience). Panel 2 included FITC rat anti-mouse CD4 (#561828, clone: GK1.5, BD Biosciences), PE-Cy7 rat anti-mouse CD45R/B220 (#561881, clone: RA3-6B2, BD Biosciences), BV421 rat anti-mouse CD279 (PD-1) (#569780, clone: RMP1-30, BD Biosciences), PE rat anti-mouse CD185 (CXCR5) (#561988, clone: 2G8, BD Biosciences), and Alexa Fluor 647 mouse anti-Bcl-6 (#561525, clone: K112-91, BD Biosciences). Data were acquired on an LSR Fortessa X20 (BD Biosciences, Franklin Lakes, NJ) or Attune NxT (Thermo Fisher Scientific, Waltham, MA) and analyzed using FlowJo_v10.10.0_CL (BD Life Sciences, Ashland, OR, USA). Doublets and debris were excluded based on FSC-A/FSC-H parameters, single color controls were used for compensation and gating purposes, and populations were defined as follows: GC B cells (CD4^-^B220^+^CD38^-^GL7^+^), plasmablasts (CD4^-^B220^+^CD138^+^), plasma cells (CD4^-^B220^-^CD138^+^), memory B cells (CD4^-^B220^+^GL7^-^CD38^+^IgD^-^IgM^-^), T helper cells (B220^-^CD4^+^), and Tfh cells (B220^-^CD4^+^CXCR5^+^PD1^+^BCL6^+^).

### Statistical analysis

Data were analyzed, and figures were generated using GraphPad Prism version 10.5.0 (GraphPad Software, CA, USA) and Microsoft Excel for Microsoft 365 (version 2507). For GTT assay, readings above the detectable threshold were assigned the maximum detectable value (i.e., 750mg/dL). To enable statistical comparisons, if measurements were below the limit of detection in antibody and TCID_50_ assays, a value that is half of the detection limit was assigned. For the cytokines/chemokines analysis, concentrations (pg/mL) were calculated using a five-parameter best line fit of the standards and samples below the detection limit were assigned half of the lowest detected value and samples above the detection limit were assigned 1.5 times the highest detectable concentration (22). Antibody titers were log_2_-transformed while cytokine concentrations were log_10-_transformed before statistical analysis.

The effects of biological sex and obesity on body mass, BMI, blood glucose levels, antibody titers, cytokines/chemokines, and B and T cell frequencies were compared using two-way ANOVA followed by Tukey’s or Bonferroni’s multiple comparisons. If repeated measurements were taken from the same animal, such as body mass change after diet-treatment or following virus challenge, and GTT, they were compared using repeated measures ANOVA (mixed effects model) with Tukey’s multiple comparisons. For comparisons between two groups such as unvaccinated vs vaccinated mice or obese versus non-responder females, the Mann-Whitney test (non-parametric) or unpaired T-test (parametric) was used. Cytokine/chemokine analysis between two groups was performed using unpaired T-test with Holm-Sidak correction for multiple comparisons. Data are shown as mean ± standard error of the mean (SEM) and considered statistically significant at p<0.05 and having a trend at 0.05≤p≤0.1.

## Results

### Diet treatment for 12 weeks induced obesity in all males but 2/3^rd^ of the female mice

Before the diet treatment began, males had significantly higher body mass than females, but there was no difference in body mass between the males or females on the two diets (**Figure 1A**). After the diet treatment, the absolute body mass of males on HFD was significantly greater from 3 weeks, while that was substantially greater in females on HFD only after 5 weeks compared to their respective controls on LFD (**Figure 1B**). After 12 weeks of diet treatment, all HFD-fed males and two-thirds of HFD-fed females developed obesity (≥20% higher body mass than sex-matched LFD controls) (**Figure 1C**). This is consistent with prior studies which showed that female mice are less responsive to obesity development (24–26). At week 12, males and females with obesity, had significantly higher body mass than their non-obese counterparts (**Figure 1D**). At 0 dpc, BMI was also elevated significantly in both groups (**Figure 1E**).

**Figure 1:**
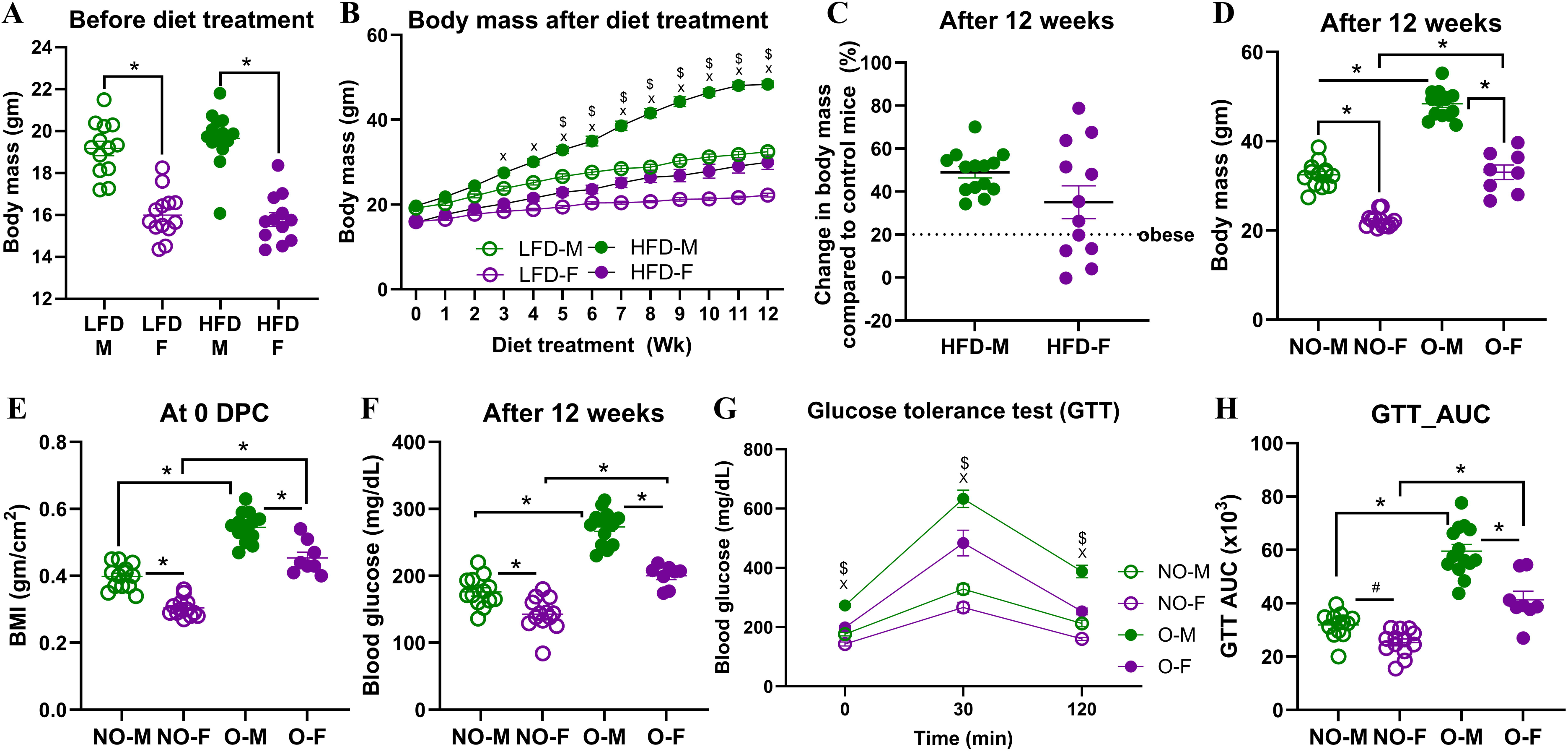
Progression of high-fat diet-induced obesity in male and female mice. Male and female C57BL/6J mice (∼5-6 weeks) were assigned to a low-fat diet (LFD) or a high-fat diet (HFD) for 12 weeks. (A) Baseline body mass before diet treatment, (B) change in body mass over 12 weeks, and (C) percent body mass increase at week 12 relative to sex-matched LFD controls are shown (n=12-14/group). Obesity was defined as ≥20% higher body mass than sex-matched LFD averages. After week 12, comparisons between non-obese and mice with obesity are shown for (D) body mass, (E) body mass index (BMI), (F) fasting blood glucose levels, (G) blood glucose excursions on glucose tolerance test (GTT) time course (0-120 min), and (H) GTT area under the curve (AUC) (n=8-14/group). Statistical comparisons were made using two-way analysis of variance (ANOVA) (A, D, E, F, H) and two-way repeated measures ANOVA (B, G) followed by Tukey’s multiple comparisons. An asterisk (*) represents a significant difference (p<0.05) and a hash (^#^) represents a trend (0.05≤p≤0.1). In figures B and G, ‘X’ and *‘*$*’* represent a significant difference (p<0.5) between the male groups and female groups, respectively. Abbreviations: males on low-fat diet (LFD-M), females on low-fat diet (LFD-F), males on high-fat diet (HFD-M), females on high-fat diet (HFD-F), non-obese males (NO-M), non-obese females (NO-F), males with obesity (O-M), and females with obesity (O-F).

Glucose tolerance testing in week 12 showed impaired fasting blood glucose and elevated glucose excursions in both groups with obesity, with significantly greater AUC values, compared to non-obese mice (**Figures 1F-H**). Prior studies have shown glucose intolerance during obesity in humans as well as in mouse models (27, 28). There was a significant effect (p<0.05) biological sex, obesity, and their interaction on body mass on 12**^th^** week of diet treatment, fasting blood glucose, and GTT AUC (**Figures 1D,F,H**). The significant effect of interaction between obesity and sex indicates the differences in response to HFD-induced obesity development in male and female mice. Yet, both males and females with obesity had higher body mass, BMI, and glucose intolerance compared to their sex-matched non-obese counterparts, confirming successful establishment of obese models in both sexes.

### Males with obesity have the lowest antibody responses following influenza vaccination

Mice were vaccinated twice with an inactivated 2009 pandemic H1N1 vaccine at a 3-week interval, and plasma samples were collected at 35 dpv to measure antibody responses. Non-obese females had significantly higher IgG and IgG2c antibodies and a higher trend of nAb titers compared to non-obese males (**Figures 2A-E**). In a similar way, female mice with obesity also had significantly higher IgG, IgG1, IgG2c, and nAb titers compared to the males with obesity (**Figures 2A-E**). There was no difference in antibody responses between non-obese females and females with obesity. However, in males, those with obesity had a lower trend of IgG2c antibody response compared to those without obesity (**Figure 2C**). Among the four groups, males with obesity had the lowest levels of antibodies, as indicated further by the higher percentages of mice (having antibodies below the limit of detection, i.e., 8% IgG, 15% IgG1, 69% IgG2c, 23% nAb) (**Figures 2A-E**). The IgG1/IgG2c ratio was comparable among the four vaccinated groups (**Figure 2D**). There was a significant effect (p<0.05) of biological sex, but not of obesity or their interaction on IgG, IgG1, IgG2c, and nAb titers (Figure 2A-E). These data suggest that sex-based differences in influenza vaccine-induced antibody responses are observed in mice with or without obesity, with females exhibiting a higher antibody quantity and quality. Moreover, males with obesity mount the lowest levels of vaccine-induced antibodies compared to other groups.

**Figure 2:**
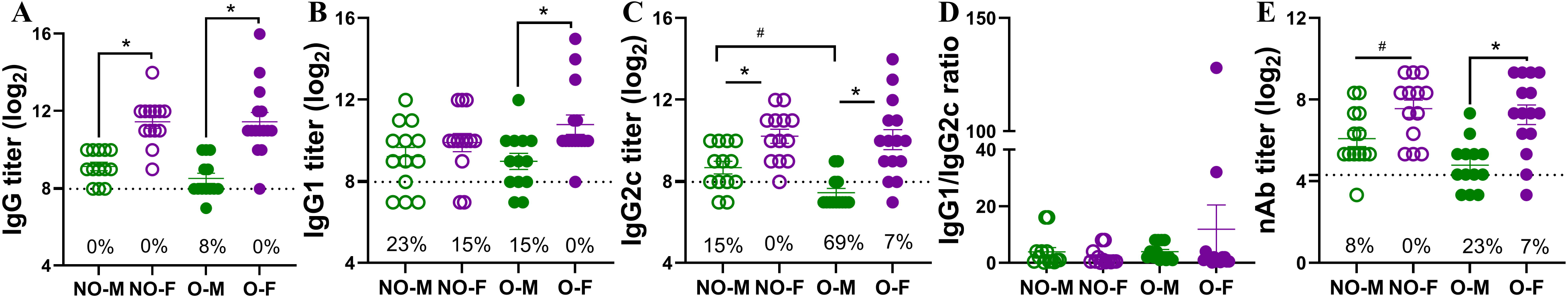
Vaccine-induced antibody responses in males and females, with or without obesity. Non-obese and obese, male/female C57BL/6J mice were vaccinated twice with the 2009 pandemic H1N1 influenza A virus (IAV) vaccine at a 3-week interval. At 35 days post-vaccination (dpv), plasma samples were collected, and antibodies were measured. (A) IgG, (B) IgG1, (C) IgG2c, (D) IgG1 to IgG2c ratio, and (E) virus-neutralizing antibody (nAb) titers are compared (dashed line – limit of detection; n=13-15/group). Subjects that had an antibody titer below the limit of detection are shown in percent in each figure. Statistical comparisons were made using two-way analysis of variance (ANOVA) (A-E) followed by Tukey’s multiple comparisons. An asterisk (*) indicates a significant difference (p<0.05) and a hash (^#^) represents a trend (0.05≤p≤0.1). Abbreviations: non-obese males (NO-M), non-obese females (NO-F), males with obesity (O-M), and females with obesity (O-F).

### Vaccinated male mice with obesity are least protected following influenza virus challenge

Vaccinated mice were challenged with a drift variant of the 2009 H1N1 virus at 42 dpv. After the virus challenge, body mass was measured for 21 dpc. The absolute (i.e., in grams) and relative (i.e., in percentage) body mass changes from 0 dpc were then compared (**Figures 3A-B**). Sex difference was observed, in mice with or without obesity, where vaccinated males suffered with greater body mass loss compared to vaccinated females. Both the absolute and relative body mass loss in non-obese males was significantly greater than non-obese females on 8 and 20 dpc. Likewise, males with obesity had significantly higher absolute (from 6 to 21 dpc) and relative (on 7, 8, 9, 10, 15, and 17 dpc) body mass loss compared to the females with obesity. The effect of obesity was also evident on body mass loss for both sexes, where vaccinated mice with obesity lost greater body mass than vaccinated non-obese mice. The females with obesity lost significantly more absolute (from 7 to 12 dpc) and relative (on 8 dpc) body mass compared to non-obese females. Likewise, males with obesity lost significantly greater absolute (from 5 to 19 dpc) and relative (from 6 to 12 dpc) body mass compared to non-obese males (**Figures 3A-B**). Subsets of vaccinated and challenged mice were euthanized at 3 dpc to determine virus titers in the lungs. While 75% of vaccinated non-obese males cleared the virus from the lungs, one tested positive (titer 4.7 log_10_ TCID_50_). The replicating virus was not detected in the lungs of any of the vaccinated non-obese females indicating a 100% virus clearance. Likewise, 75% of vaccinated females with obesity cleared the virus from the lungs, and one tested positive with a low virus titer (i.e., 2.3 log_10_ TCID_50_). Remarkably, 100% (4/4) of the vaccinated males with obesity tested positive with a high-replicating virus titer (3.9 to 5.4 log_10_ TCID_50_) in the lungs, which was significantly higher than in females with or without obesity (**Figure 3C**).

**Figure 3:**
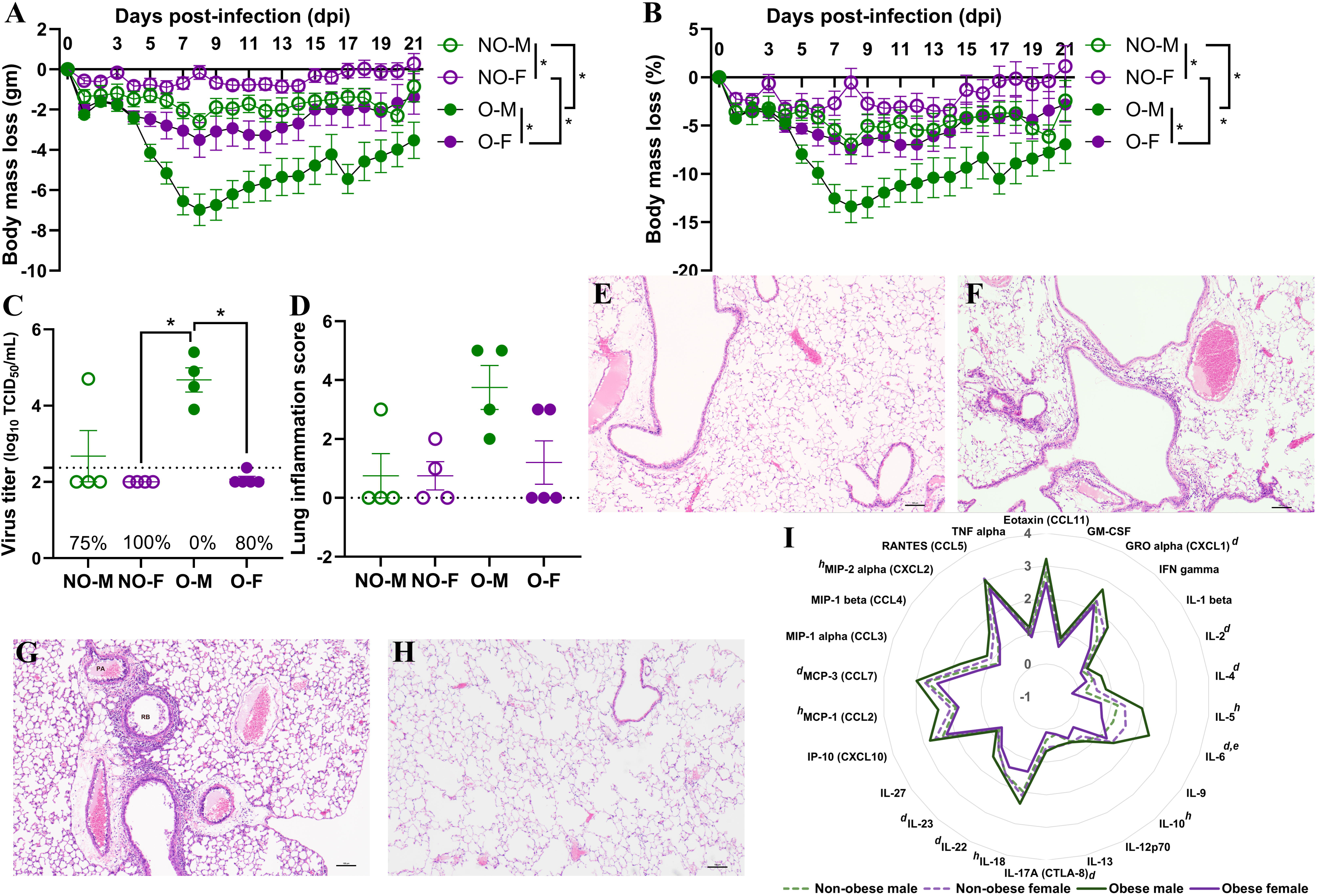
Post-challenge morbidity, virus replication, and inflammatory changes in vaccinated mice. Vaccinated non-obese and obese, male/female mice were challenged with a drift variant of the 2009 H1N1 IAV at 42 days post-vaccination (dpv). (A) Absolute body mass change (g), and (B) relative change (%) through 21 days post-challenge (dpc) is shown (n=5-6/group). (C) Lung virus titers at 3 dpc (TCID_50_ assay, dashed line: limit of detection) and (D) lung inflammation scores at 3 dpc (H&E staining) are compared (n=4-5/group). Representative H&E images of lungs at 3 dpc from a (E) vaccinated non-obese male, (F) non-obese female, (G) males with obesity, and (H) females with obesity are presented. (I) Radar plot of log_10_-transformed cytokines/chemokines concentrations in lung homogenates at 3 dpc are compared (n=3-5/group). Statistical comparisons were made using (A-B) two-way repeated measures analysis of variance (ANOVA) followed by Tukey’s multiple comparisons, (C, D) Kruskal-Wallis test followed by Dunn’s post-hoc comparisons, or (I) two-way ANOVA followed by Bonferroni’s multiple comparisons. An asterisk (*) indicates a significant difference (p<0.05) and a hash (^#^) represents a trend (0.05≤p≤0.1). In Figure I, *a* or *e* respectively represent a significant difference or trend between males with or without obesity, *b* or *f* between females with or without obesity, *c* or *g* between non-obese males and non-obese females, and *d* or *h* between males and females with obesity. Abbreviations: non-obese males (NO-M), non-obese females (NO-F), males with obesity (O-M), and females with obesity (O-F).

Pulmonary inflammation was evaluated at 3 dpc using histopathology and measurement of various cytokines/chemokines. Inflammatory changes were not observed in majority of vaccinated non-obese males, non-obese females, and females with obesity. Lung inflammation was evident in all of the vaccinated males with obesity and inflammation scores were higher than all other groups, however, data were not statistically significant (p>0.05) (**Figure 3D**). Representative images of H&E-stained lung tissues from non-obese males, non-obese females, males with obesity, and females with obesity are shown, respectively (**Figure 3E-H**). In accordance, the overall cytokine/chemokine concentrations in the lungs at 3 dpc were higher in males with obesity compared to other groups (**Figure 3I**). Statistically, the concentrations of GROα, IL-2, IL-4, IL-6, IL-17A, IL-22, IL-23, and MCP-3 were significantly higher in males with obesity compared to the females with obesity, while the concentrations of IL-5, IL-10, IL-18, MCP-1, and MIP2α trended higher (*p*≤*0.1*) (**Figure 3I**). Males with obesity also had a higher trend (*p*≤*0.*1) for the concentration of IL-6 compared to non-obese males. These data indicate that vaccinated male mice with obesity were least protected compared to other vaccinated groups, as evidenced by higher body mass loss, inability to clear viruses from the lungs, and higher concentrations of inflammatory cytokines and chemokines.

### Vaccination is still beneficial to all groups, including males with obesity

To determine if vaccination was beneficial in all groups, including the males with obesity, mock-vaccinated but virus-challenged mice were used as controls and euthanized at 3 dpc to compare pulmonary virus replication, histopathological changes, and cytokines/chemokines. In all groups, unvaccinated mice had virus titers ranging between 4.7 to 6.5 log_10_ TCID_50_ (**Figures 4A-D**). Compared to unvaccinated mice, vaccinated non-obese males, vaccinated males with obesity, vaccinated non-obese females, as well as vaccinated females with obesity, all had significantly lower virus titers in the lungs (**Figures 4A-D**). Histopathology analysis also showed a significant reduction in the inflammation in the lungs of vaccinated mice compared to unvaccinated mice, for all groups (**Figures 4E-H**). Likewise, when the cytokine/chemokine responses were compared in the lung homogenates of unvaccinated versus vaccinated and virus-challenged mice, vaccinated mice had overall lower concentrations of most of the inflammatory cytokines/chemokines tested (**Figures 4I-L**). Specifically, in non-obese males, concentration of GRO-α was significantly reduced (p<0.05) and that of IL-6 and IL-23 showed a lower trend in vaccinated mice (**Figure 4I**). In non-obese females, concentrations of GM-CSF, GRO-α, IL-1β, IL-6, IP-10, MCP-1, MCP-3, MIP-1α, MIP-2α, and TNF-α were significantly lower and eotaxin has a lower trend in vaccinated mice of compared to their unvaccinated controls (**Figure 4J**). In males with obesity (**Figure 4K**), concentration of GRO-α was significantly lower (p<0.05) and IL-6 showed a lower trend in vaccinated mice. Likewise, in females with obesity (**Figure 4L**), vaccinated mice had significantly lower levels of eotaxin, GM-CSF, GRO-α, IL-1β, IL-2, IL-5, IL-6, IL-12p70, IL-17A, IL-18, IP-10, MCP-1, MCP-2, MIP-1α, MIP-1β, MIP-2α, and TNF-α compared to unvaccinated controls. The differences in numbers of cytokines and chemokines showing significant differences in males versus females, irrespective of obesity, likely indicates the development of severe disease in unvaccinated females than in males during influenza virus challenge as shown earlier (22, 29). These data indicate that, though the vaccine was less effective in males with obesity, it still protected them by significantly reducing the virus replication and inflammation in the lungs.

**Figure 4:**
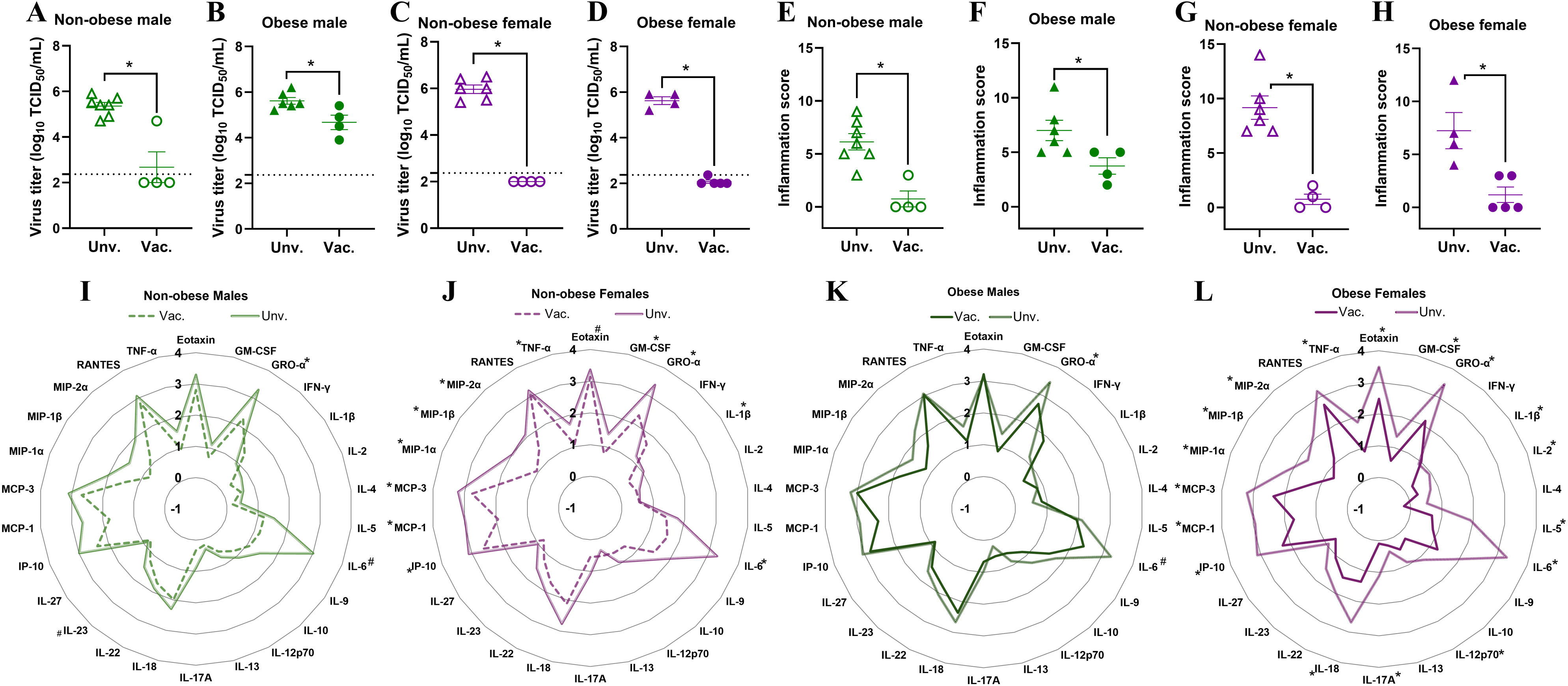
Comparisons of virus replication and inflammatory changes between unvaccinated (unv.) and vaccinated (vac.) mice. Non-obese and obese, male/female mice were either vaccinated (prime-boost) or vehicle-treated and challenged with 10^5^ TCID_50_ units of drifted 2009 pandemic H1N1 influenza A virus (IAV). At 3 days post-challenge (dpc): (A-D) lung viral titers (dashed line: limit of detection) and (E-H) inflammation scores are shown for (A, E) non-obese males, (B, F) non-obese females, (C, G) males with obesity, and (D, H) females with obesity (n=4-7/group). (I-L) Radar plots show log_10_-transformed cytokine/chemokine concentrations comparing unvaccinated vs vaccinated mice within each group (n=3-7/group). Statistical comparisons were made using (A-H) Mann-Whitney test and (I-L) unpaired T-test followed by Holm-Sidak test for multiple comparisons. An asterisk (*) indicates a significant difference (p<0.05) and a hash (^#^) represents a trend (0.05≤p≤0.1).

### Splenic B cell responses are comparable among vaccinated mice while plasma cells in bone marrow are lower in males, irrespective of obesity status

To test if the lower antibody responses and inferior protection in males with obesity was associated with a lower frequency of B and Tfh cells, we collected spleen from vaccinated mice at 35 dpv and performed flow cytometry to quantify the frequencies of germinal center B cells, plasmablasts, plasma cells, memory B cells, and Tfh cells (**Supplementary Figures 1-2**). Likewise, bone marrow cells were also collected at 35 dpv to test if the frequencies of B cells and plasma cells differ in bone marrow (**Supplementary Figure 3**). In the spleen, the frequencies of CD4^-^B220^+^ B cells were comparable among the four vaccinated groups (**Figure 5A**). Two-way ANOVA analysis showed the effect of obesity on the germinal center (CD4^-^B220^+^CD38^-^GL7^+^) B cells, which were significantly higher in mice with obesity compared to non-obese mice (**Figure 5B**). The frequencies of plasmablasts, plasma cells, and memory B cells, however, were not-significantly different based on biological sex and obesity status (**Figures 5C-E**). The frequencies of T helper (B220^-^CD4^+^) cells and Tfh cells in spleen were also comparable among the four vaccinated groups (**Figures 5F-G**). In the bone marrow, while the B cell frequencies were comparable, the frequencies of plasma cells were significantly higher in non-obese females compared to non-obese males (**Figures 5H-I**). Similar effect of biological sex was also observed in mice with obesity, as vaccinated females with obesity had a higher trend (*p=0.05*) of plasma cells in the bone marrow compared to the males with obesity (**Figure 5I**).

**Figure 5:**
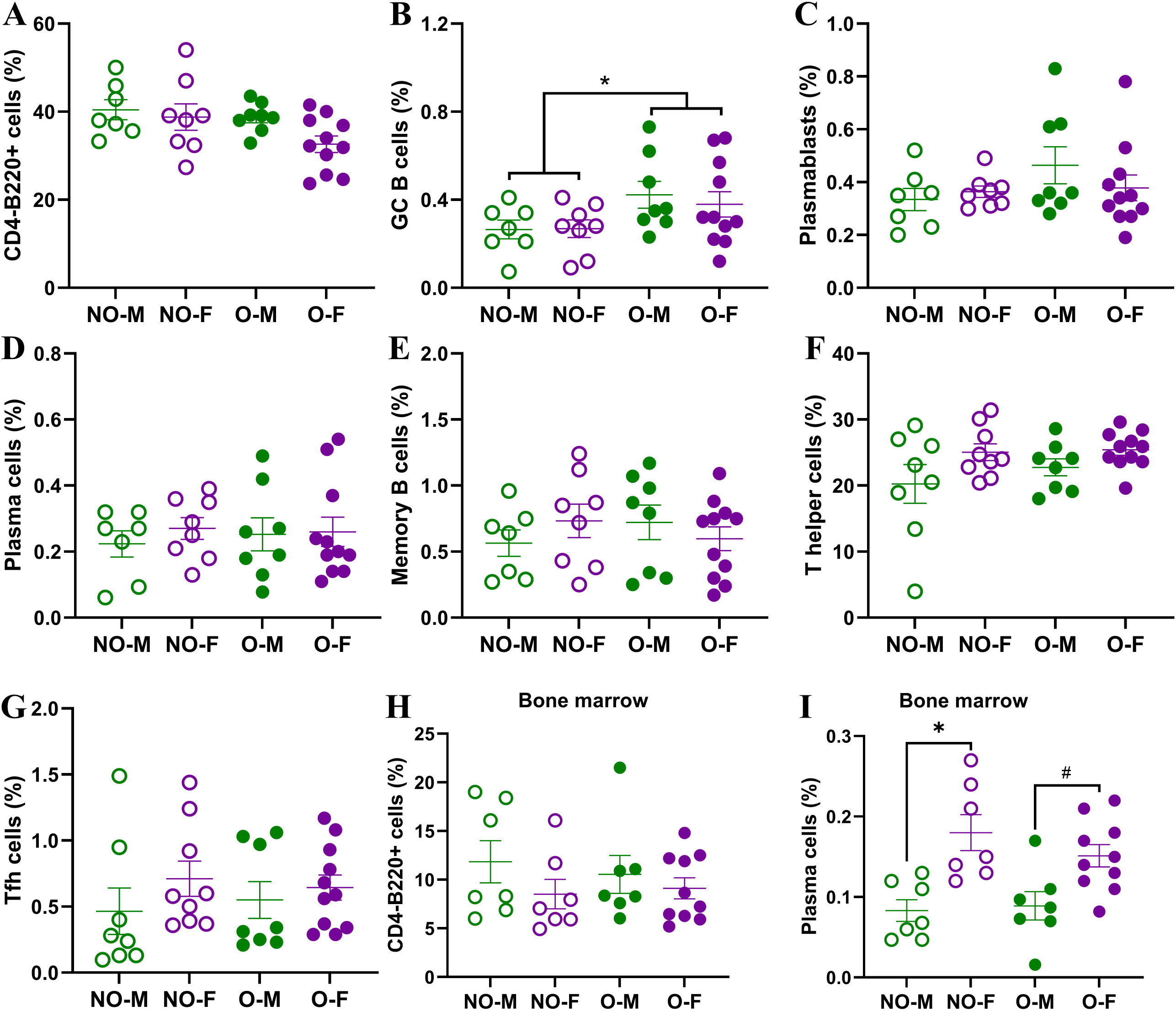
B- and T-cell responses in spleen and bone marrow of vaccinated mice. At 35 days post-vaccination (dpv) flow cytometry was performed on splenic and bone marrow cells to determine frequencies of various B and T cells. In spleen, the frequencies of (A) CD4^-^B220^+^ B cells, (B) CD4^-^B220^+^CD38^-^GL7^+^ germinal center (GC) B cells, (C) CD4^-^B220^+^CD138^+^ plasmablasts, (D) CD4^-^B220^-^CD138^+^ plasma cells, (E) CD4^-^B220^+^GL7^-^CD38^+^IgD^-^IgM^-^ memory B cells, (F) B220^-^CD4^+^ T helper cells, and (G) B220^-^CD4^+^CXCR5^+^PD1^+^BCL6^+^ T follicular helper (Tfh) cells were quantified (n=7-11/group). In bone marrow, frequencies of (H) CD4^-^B220^+^ B cells, and (I) CD4^-^B220^-^CD138^+^ plasma cells were quantified (n=7-10/group). Statistical comparisons were made using two-way analysis of variance (ANOVA), and an asterisk (*) indicates a significant difference (p<0.05) and a hash (^#^) represents a trend (0.05≤p≤0.1). Abbreviations: non-obese males (NO-M), non-obese females (NO-F), males with obesity (O-M), and females with obesity (O-F).

### Females on HFD, responders and non-responders, have comparable vaccine-induced immunity and protection

As nearly 30% of female mice on HFD were non-responders, i.e., they did not become obese even after being on HFD, we aimed to determine the differences in vaccine-induced immunity and protection between the non-responder and responder (i.e., the obese) females (**Figure 6**). Before the diet treatment started, the body mass of mice in both groups was comparable (**Figure 6A**). Following the diet treatment, the difference in body mass was evident from the 3^rd^ week onwards (**Figure 6B**). The females with obesity had significantly higher body mass and BMI (**Figure 6C-D**). Interestingly, despite having significantly higher body mass, the blood glucose levels over time on GTT and GTT AUC were comparable between obese and non-responder females (**Figures 6E-F**). The IgG, IgG1, IgG2c antibody titers, IgG1/IgG2c antibody ratios, and nAb titers were also comparable at 35 dpv between obese and non-responder females (**Figures 6G-K**).

**Figure 6:**
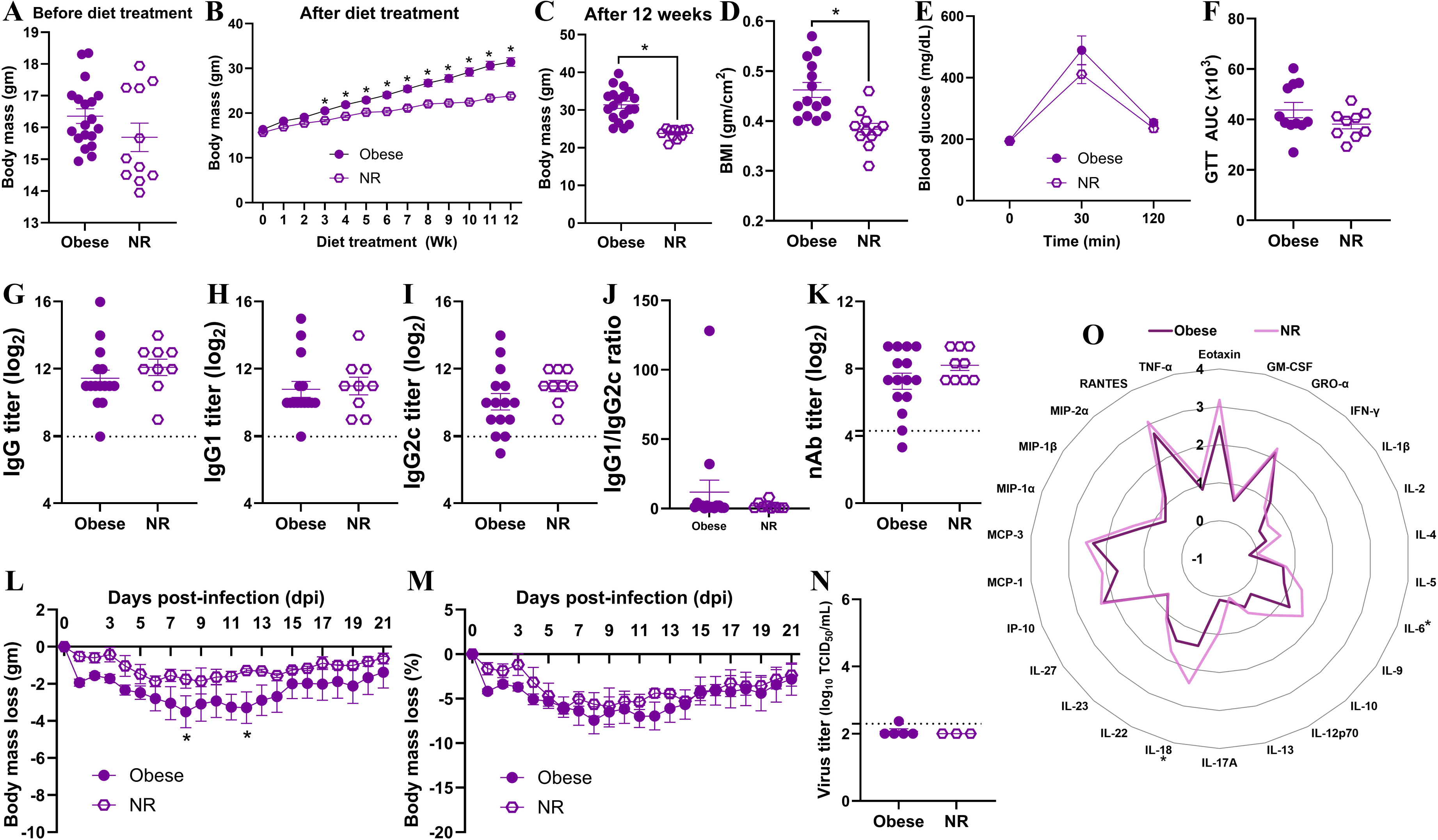
Comparisons of immunity and protection between non-responder (i.e., NR) and responder (i.e., obese) females on high-fat diet. Following 12 weeks on HFD, nearly 2/3^rd^ of females developed obesity, and the remainder were non-responders. (A) Baseline body mass; (B) body mass over 12 weeks on HFD; (C) week 12 body mass; (D) body mass index (BMI); (E) glucose tolerance test (GTT) time course (0 – 120 min); and (F) GTT area under the curve (AUC) are shown (n=9-19/group). After prime-boost vaccination with the inactivated 2009 H1N1 influenza vaccine, at 35 days post-vaccination (dpv), (G) IgG, (H) IgG1, (I) IgG2c, (J) IgG1/IgG2c, and (K) virus-neutralizing antibody (nAb) titers were measured (dashed line – limit of detection, n=9-15/group). Following the challenge with the drift variant of H1N1 virus, (L) absolute (g) and (M) relative (%) body mass change through 21 days post-challenge (dpc); (N) lung viral titers at 3 dpc; and (O) log_10_-transformed cytokines/chemokines concentrations at 3 dpc were compared (n=3-5/group). Statistical comparisons were made using the (A, C-K) Mann-Whitney test, (B, L, M) two-way repeated measures analysis of variance (ANOVA) followed by Tukey’s multiple comparisons, and unpaired T-test followed by Holm-Sidak test for multiple comparisons. An asterisk (*) indicates a significant difference (p<0.05) and a hash (^#^) represents a trend (0.05≤p≤0.1).

After the virus challenge, the absolute body mass change among females with obesity was significantly higher on 8^th^ and 12^th^ dpc, while no such difference was observed on the percentage change in body mass (**Figures 6L-M**). Likewise, when replicating virus titers were compared at 3 dpc in the lung homogenates, they were comparable between non-responders and females with obesity (**Figure 6N**). When cytokines/chemokines were compared, vaccinated non-responder females had significantly higher concentrations of IL-6 and IL-18, while other cytokines/chemokines were comparable (**Figure 6O**). Histopathology analysis of lung tissues also showed no significant differences (data not shown). These data suggest that, despite some differences in cytokine induction, there were no major differences in antibody responses and protection between the females on HFD regardless of whether they developed obesity or not.

## Discussion

Using the diet-induced obesity mouse model, we investigated whether sex difference in influenza virus vaccine responses is observed during obesity or not. Our findings showed that the effects of biological sex on influenza vaccine-induced immunity and protection exists even during obesity with a better response driven by female sex. Specifically, vaccinated male mice with obesity generated the weakest humoral immune response, had more severe disease, were unable to clear viruses from the lungs, and suffered from more severe pulmonary inflammation despite influenza virus vaccination. However, female mice, irrespective of diet and obesity status, developed higher antibody response and were better protected.

Following vaccination with the 2009 pandemic H1N1 IAV vaccine, female non-obese mice had higher induction of antibodies and were better protected, compared to the non-obese males. Our finding that non-obese females mount higher antibody responses and are better protected than males, following influenza vaccination, is consistent with previous studies both in humans and mouse models (14–16, 19, 21, 22, 30). In non-obese mouse models, the female-biased higher antibody response and better protection persist up to 4 months post-vaccination (21), and extends to diverse IAV mutants (19). Robust responses to influenza vaccine in non-obese females are associated with higher *Aicda* gene expression and somatic hypermutation (19); and increased expression of the X-linked *Tlr7* gene in the splenic B cells, regulated by increased DNA methylation (17). Moreover, such sex-biased differences in influenza vaccine efficacy are primarily driven by sex steroids (15, 16), more than chromosomes, as shown by studies involving gonadectomy and exogenous hormone replenishment, and the four-core genotype (FCG) mouse model (15, 21).

Our study showed that sex difference exists in influenza vaccine efficacy even during obesity, at least in a mouse model. Males with obesity produced fewer antibodies, failed to clear the virus from the lungs, and had exaggerated lung inflammation. Vaccinated males with obesity had elevated levels of proinflammatory cytokines and chemokines including IL-6, the central mediator of acute inflammation; GRO-α, MIP-2α, and IL-17A, which are potent neutrophil chemoattractant that also promote lung inflammation; and MCP-1 and MCP-3, which recruit monocytes and other leucocytes and contribute to lung damage (31, 32). Even in females with obesity, despite having similar antibody responses to non-obese females, they suffered with more severe disease, as shown by significantly greater body mass loss and failure to recover to baseline body mass. Human studies parallel this, showing that obesity accelerates antibody decline and increases the risk of influenza-like illness (7, 8). Prior studies suggest that obesity-associated dysfunctions during influenza vaccination include reduced switched memory and transitional B cells, impaired CD8^+^ T-cell activation and function, and altered cytokine production (7, 33, 34). Importantly, weight loss or dietary changes before vaccination can improve influenza vaccine-induced immunity and protection in both males and females with obesity (33, 35).

Although vaccine efficacy was reduced in males with obesity, vaccination still offered protection, as evidenced by reduced lung virus titers and inflammatory cytokines compared to the unvaccinated controls. This is consistent with previous findings in humans and in mouse models that vaccination is still beneficial to all, including males with obesity, to protect them from influenza-associated severe disease (8–10). Therefore, the current vaccination scheme should be continued while exploring novel ways to improve influenza vaccines for the population with obesity.

Non-responder females, which remained non-obese despite the HFD-treatment, developed antibody responses comparable to females with obesity. In a prior study, intraperitoneal immunization with the A/Puerto Rico/8/34 vaccine resulted in no significant difference in antibody titers between the non-responder and responder females; however, the absolute body mass loss was significantly higher in females with obesity (36). We also observed significantly greater absolute body mass loss in females with obesity on certain days, and there were subtle cytokine/chemokine differences in the lungs. However, considering no such differences in relative body mass loss, virus clearance from the lungs, and pulmonary inflammation, we conclude that there was no difference in influenza vaccine response between responder and non-responder females on HFD. Therefore, overall greater immunity in females following influenza vaccination is conserved across different metabolic states, non-obese, non-responder in HFD, or with obesity. This is likely to be mediated by conserved female sex-driven immune advantages, such as circulating estradiol, which is known to activate and promote the survival of B cells (21, 37, 38).

Vaccine studies that stratified data by biological sex and obesity are limited. Yet, following COVID-19 vaccination, higher BMI was associated with lower titers of antibodies in males, but not in females (39, 40). A recent study using a mouse model of DIO also showed that following mRNA-1273 vaccination, females with obesity produced significantly higher antibodies compared to the males (41). In our study, despite comparable splenic B- and Tfh-cells in males and females with obesity, differences in antibody responses and protection were observed. This suggests that B-cell antibody production following influenza vaccination was more efficient in females than in males with obesity. The higher frequency of bone marrow plasma cells in females may contribute to their superior antibody response. To investigate the hypothesis of sex-specific impairment in antibody production by B cells (or B-cell inefficiency) during obesity, future research should investigate sex-specific differences in B-cell cellular metabolism, mitochondrial functions, metabolomics, and somatic hypermutation.

Sex steroids have a significant role in shaping immune responses to influenza vaccines (15, 16, 19, 21). Therefore, further work is necessary to determine how sex steroids and obesity affect B-cell function and influenza vaccine responses, particularly exploring the sex-specific impairments observed in males. Estrogen enhances B-cell differentiation, germinal center formation, and influences cellular metabolic functions (21, 42, 43). In non-obese humans, estradiol is positively correlated with influenza vaccine-induced antibody levels, and testosterone is negatively correlated (15, 16). Estradiol has a pro-inflammatory response and activates B cells, while testosterone has an anti-inflammatory response and suppresses the B cells (37, 38). Estradiol upregulates genes like *Bcl-2,* which promotes the survival of B cells, increases the production of B-cell activation factor (BAFF), and induces polyclonal activation of B cells, while testosterone negatively regulates these changes, and for this reason, autoimmunity is more prevalent in females than in males, and increases in males with testosterone deficiency (44–46). Approaches such as the use of a four-core genotype (FCG) mouse model to segregate the effects of gonadal hormones or sex chromosomes (21), and gonadectomy and hormone replacement (15, 22), will help in understanding the direct role of sex steroids in mediating sex differences in influenza vaccine-induced immunity and protection in mice with obesity.

Mouse models have limitations, as they do not recapitulate every aspect of human obesity. The C57BL/6J mice are inbred with limited genetic and metabolic heterogeneity and develop obesity over a shorter period than humans. In our study, females with obesity remained smaller and less hyperglycemic than males with obesity, potentially underestimating sex-specific effects, and the role of hyperglycemia warrants further investigation. The mouse model we used is a naïve mouse model that lacks prior immunity, whereas most humans are primed by prior infection or vaccination. Future studies could employ outbred mouse populations such as the collaborative cross strains to better capture genetic diversity, use thermoneutral housing to induce severe obesity and hyperglycemia in females, and incorporate prior-infection model to represent human immune history. Likewise, we tested a monovalent H1N1 vaccine and the results may not generalize to seasonal influenza virus vaccines that are either trivalent or quadrivalent. Therefore, vaccine studies with other strains and subtypes, influenza B viruses (IBVs), and seasonal tri- or quadrivalent vaccines, and subsequent challenge experiments with diverse strains and subtypes will be required to obtain a broader perspective on the interaction between obesity and biological sex in influenza vaccine efficacy.

Our study provides novel insights into how the dynamic interactions of biological sex and obesity influence the outcome of influenza vaccination. The effect of obesity on influenza vaccine efficacy is more severe in males than in females. In females with obesity, there is preservation of vaccine-induced immunity, reflecting mechanisms that may be contributed by sex hormones and X-linked immune gene expression. These findings highlight the importance of considering both sex and obesity in vaccine evaluation. Sex-inclusive studies are crucial to identify mechanisms of immune dysfunction and inspire tailored vaccine strategies. Future approaches may include adjuvants targeting B-cell metabolism, optimized dosing, or personalized vaccine schedules for individuals with obesity.

## Conflict of Interest

The authors declare that the research was conducted in the absence of any commercial or financial relationships that could be construed as a potential conflict of interest.

## Author Contributions

Conceptualization, funding acquisition, project administration, and supervision: SD; Methodology, data curation, formal analysis, validation, visualization: BW, SP, SV, SBM, TA, SD; Writing – original draft: BW, SD; Writing – review and editing: BW, SP, SV, SBM, TA, SD.

## Funding

National Institute of General Medical Sciences (NIGMS) of the National Institutes of Health (NIH) through the Center on Emerging and Zoonotic Infectious Diseases (CEZID) at Kansas State University (KSU) under the award number P20GM130448 and the start-up funds provided to S.D. by the College of Veterinary Medicine at KSU.

## Acknowledgments

The vaccine and challenge viruses were kindly provided by Drs. Sabra Klein and Andrew Pekosz (Johns Hopkins Bloomberg School of Public Health, Baltimore, MD, USA). The authors thank the Comparative Medicine Group (CMG) at Kansas State University (KSU) for their assistance in animal studies. We also acknowledge the Animal Model/Pathology (AMP) and Molecular and Cellular Biology (MCB) cores of the Center on Emerging and Zoonotic Infectious Diseases (CEZID) at KSU (supported by NIGMS award number P20GM130448) for their support during animal protocol development and flow cytometry assays.

## Data availability

Data presented in this study jwill be available upon publication and upon request from the corresponding author.

## Notes

### Competing Interest Statement

The authors have declared no competing interest.

