## Supplementary figures for "Females with Obesity Exhibit Greater Influenza Vaccine-induced Immunity and Protection than Males in a Mouse Model"

Running title: Sex Difference in Flu Vaccine Response During Obesity

Brian Wolfe[1,#], Saurav Pantha[1,#], Saranya Vijayakumar[1], Shristy Budha Magar[1], Tawfik Aboellail[1,2], Santosh Dhakal[1,*]

[1]Department of Diagnostic Medicine/Pathobiology (DMP), College of Veterinary Medicine (CVM), Kansas State University (KSU), Manhattan, KS, 66506.

[2]Kansas State Veterinary Diagnostic Laboratory (KSVDL), College of Veterinary Medicine (CVM), Kansas State University (KSU), Manhattan, KS, 66506.

[#]The authors contributed equally to this project.

*Correspondence:

Dr. Santosh Dhakal

Department of Diagnostic Medicine/Pathobiology (DMP)

Kansas State University (KSU), Manhattan, KS, 66506

**Supplementary Figure 1:**

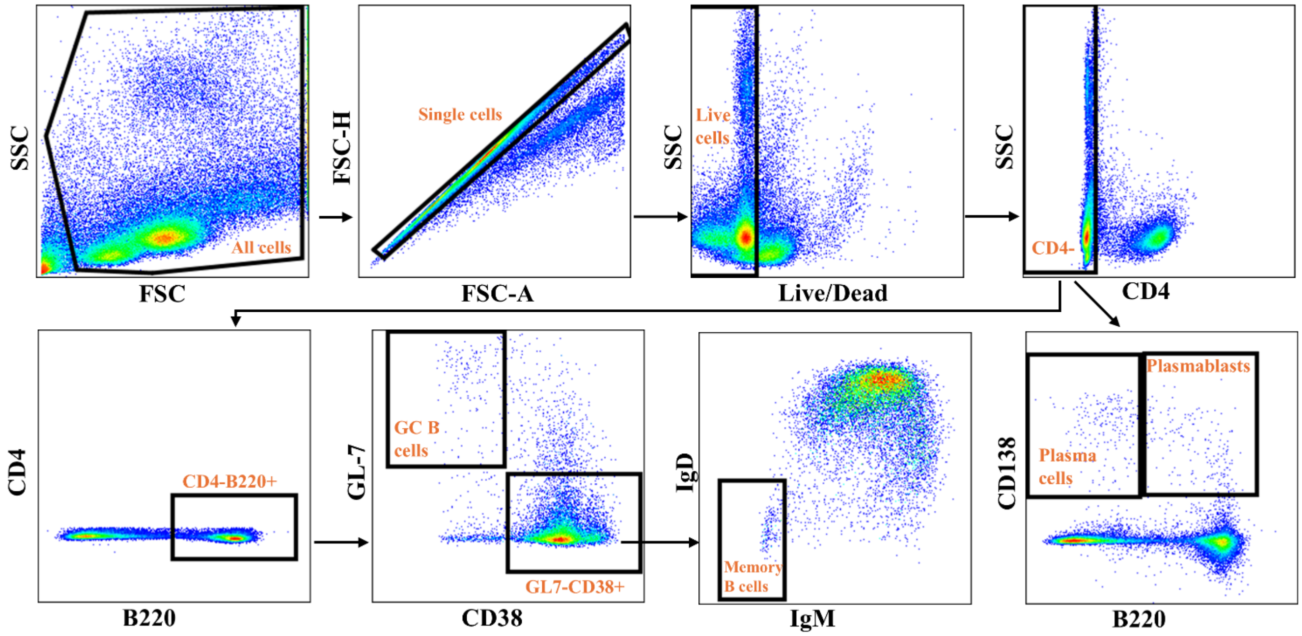


**Gating strategy for splenic B-cell subsets.** At 35 days post-vaccination, splenocytes were processed for flow cytometry, and a representative gating strategy is shown. It included: lymphocytes (FSC/SCC), single cells (FSC-H/FSC-A), live cells (viability dye negative), $CD4^-$ (to exclude T cells), followed by plasmablasts ($B220^+CD138^+$) and plasma cells ($B220^-CD138^+$). From $CD4^-B220^+$ cells: germinal center (GC) B cells ($CD38^-GL7^+$) and memory B cells ($GL7^-CD38^+IgD^-IgM^-$) were gated.

**Supplementary Figure 2**

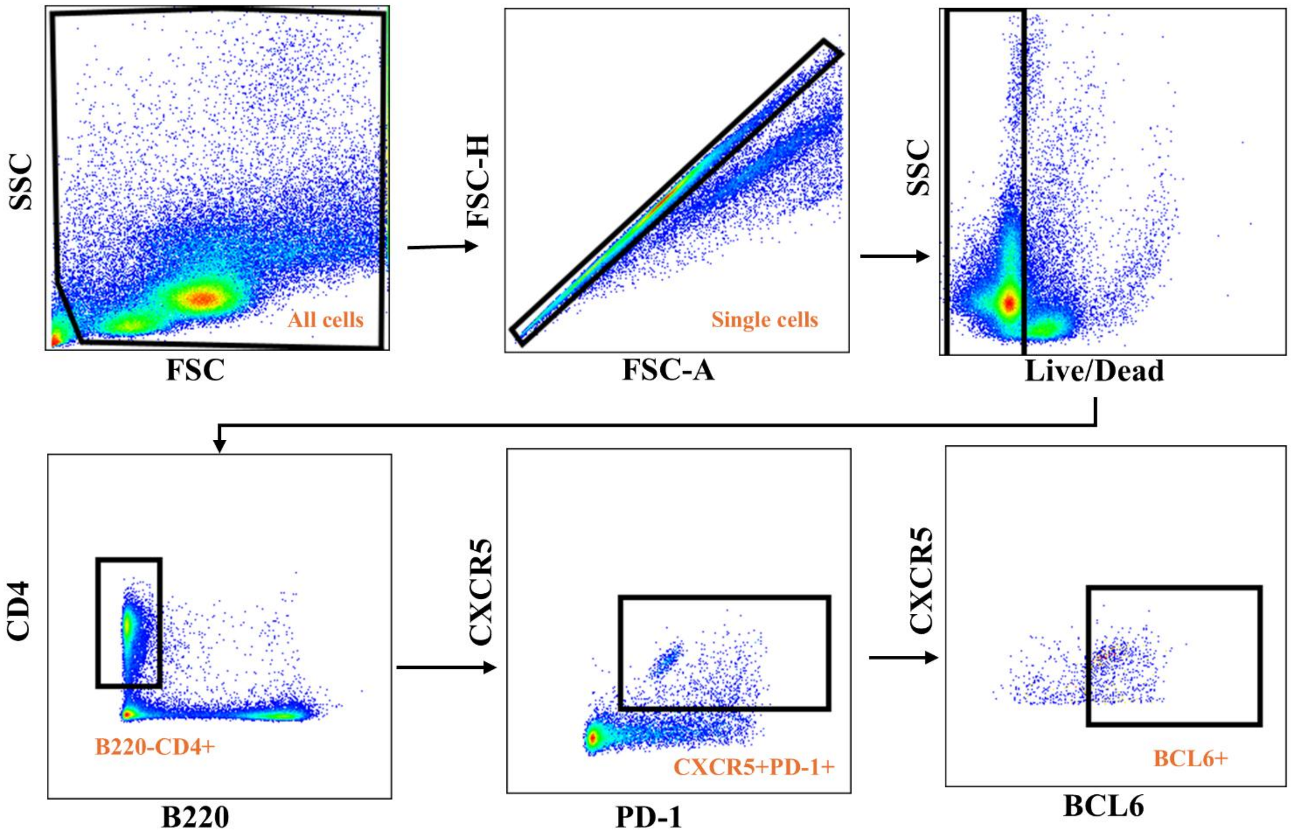


**Gating strategy for splenic T follicular helper (Tfh) cells.** Gating strategy included: lymphocytes (SSC/FSC), single cells (FSC-H/FSC-A), live cells (viability dye negative), $CD4^{+}B220^{-}$ (to exclude B cells), followed by gating for $CXCR5^{+}PD1^{+}$ cells and Tfh cells ($CXCR5^{+}PD\text{-}1^{+}BCL6^{+}$).

**Supplementary Figure 3**

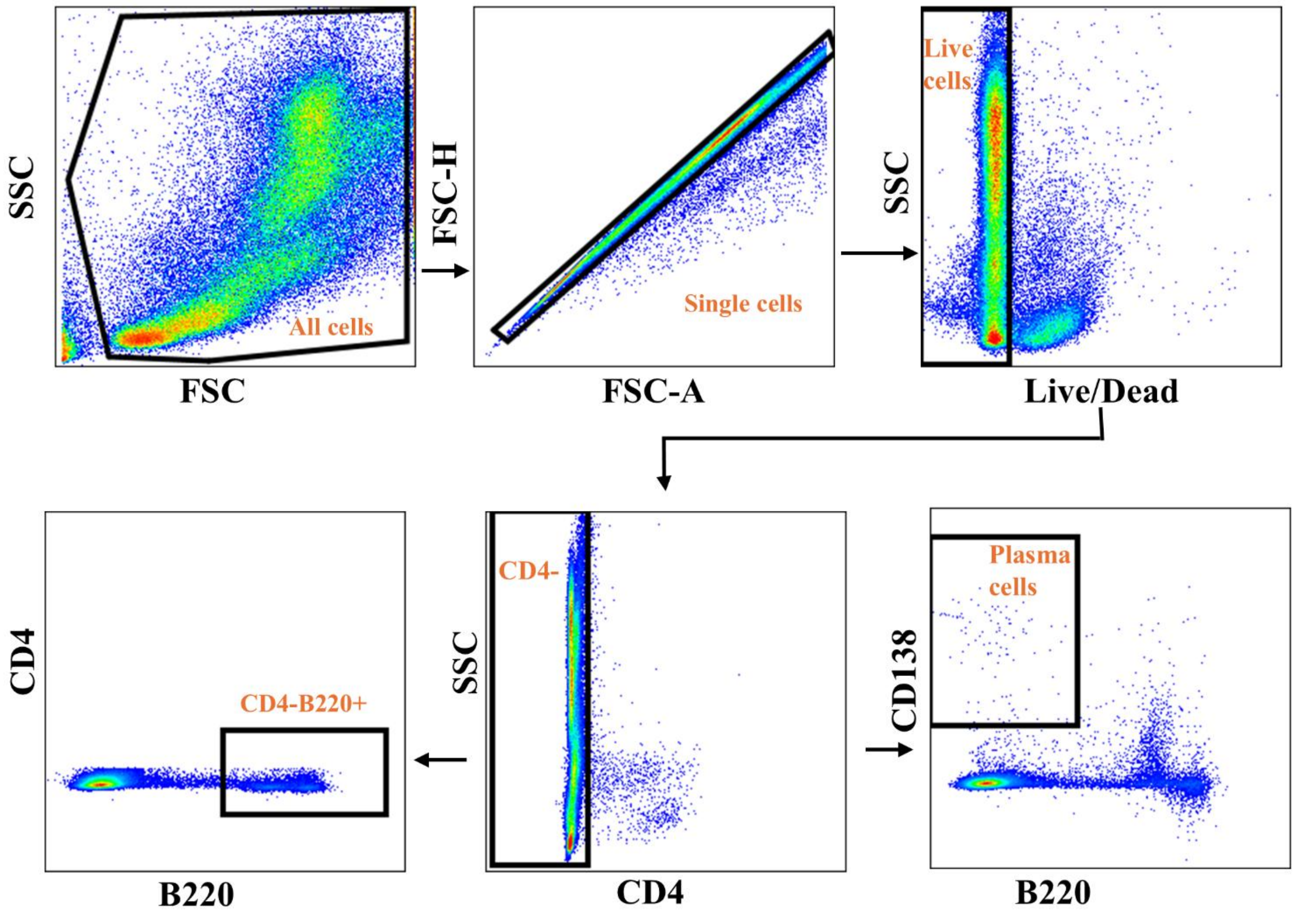


**Gating strategy for bone marrow plasma cells.** Gating strategy included: lymphocytes (SSC/FSC), single cells (FSC-H/FSC-A), live cells (viability dye negative), $CD4^-$ (to exclude T cells), followed by B cells ($CD4^-B220^+$) or plasma cells ($B220^-CD138^+$).
